# A high-performance end-to-end 3D CLEM processing workflow for facilities

**DOI:** 10.64898/2026.03.13.711046

**Authors:** Hélène Roberge, Tatiana Woller, Benjamin Pavie, Julian Hennies, Cecilia de Heus, Lakshmi Edakkandiyil, Nalan Liv, Sebastian Munck

**Affiliations:** VIB Bio Imaging Core and VIB-KU Leuven Center for Neuroscience, KULeuven Department of Neurosciences, Leuven, Belgium; VIB Bio Imaging Core and VIB-UGent Center for inflammation research, Ghent, Belgium; European Molecular Biology Laboratory (EMBL), EMBL Imaging Centre, Heidelberg, 69117, Germany; Cell Biology, Center for Molecular Medicine, University Medical Center Utrecht, Utrecht University, Utrecht, Netherlands

## Abstract

Correlative Light and Electron Microscopy (CLEM) integrates the molecular specificity of light microscopy (LM) with the ultrastructural detail of electron microscopy (EM), enabling comprehensive spatial analysis of biological samples. Despite growing demand, processing 3D CLEM datasets remains challenging, specifically for service provision in facilities, due to their multimodal nature and the lack of unified approaches. Typical steps include EM slice alignment, LM–EM registration, segmentation, and 3D visualization. We present a modular, end-to-end pipeline that consolidates existing and newly developed tools into a coherent workflow for 3D CLEM analysis and allows railroading the approach. Designed as interoperable modules accessible through a user-friendly interface, the pipeline is fully open-source and scales from standard workstations to high-performance computing environments to address the need for analysis of growing datasets. While some steps still require manual input, individual components can be automated to increase throughput and reproducibility. Together, this integrated solution lowers technical barriers and supports broader adoption of 3D CLEM methodologies.

## 1. Introduction

Correlative Light (LM) and Electron Microscopy (EM) (CLEM) workflows are powerful techniques that bridge the gap between fluorescence microscopy and ultrastructural electron microscopy^1–3^. Over the last decade, CLEM workflows were constantly improved by the implementation of live imaging^3–5^, super-resolution techniques in LM^6–8^, volume EM techniques^9–11^ such as EM tomography^12^ or 3D Focused Ion Beam/Scanning Electron Microscopy (FIB/SEM)^13,14^ and the utilisation of cryogenic conditions^7,15^. The integration of 3D LM and EM imaging techniques within CLEM workflows allows combining the molecular specificity of fluorescence microscopy with the high-resolution ultrastructural context of electron microscopy (EM) and hence further improves spatial understanding of samples, increasing their content and, consequently, scientific impact^2,9,11,16^. To stay aligned with new advancements and ensure effective image visualization and exploitation, image processing methods must also constantly be adapted^17^.

The computational analysis of the CLEM data is complex due to its multimodal nature^1^. As a result, several publications have focused on individual problems related to CLEM data. Despite the growing demand, 3D CLEM existing workflows depend on specific image processing tools and pipelines that are highly fragmented and inconsistent. Therefore, there is an unmet need for a more unified approach. In fact, due to its specialized nature and uniqueness^10,18,19^, there are many ways to execute CLEM. This could be partially explained by the multistep nature of CLEM processing, which requires precise correlation among different imaging techniques, each with its own specifications, such as pixel size, field of view, and 3D imaging. In addition, 3D CLEM workflows require a broad range of steps, such as i) EM data pre-processing with alignment and denoising (optional) ^20–22^, ii) registration between the LM and EM volume^10,18^, iii) segmentation of a region of interest^17,23,24^, and iv) finally 3D visualisation and animation^10,25–27^. To the best of our knowledge, no clear, well-defined, and freely available recipe exists to guide researchers through the many steps, from image alignment to quantitative analysis, in a modular and scalable way applicable to growing datasets.

On top of that, many labs cannot afford commercial software to perform image processing, which contributes to the fragmentation of CLEM workflows. Consequently, to promote accessibility and harmonize practices, the use of open-source software such as Napari^28^, Fiji^27,29^, Blender^30,31^ and other Python-based programs is particularly advantageous^28^. Furthermore, the large amount of data generated in CLEM applications and segmentation workflows requires new strategies for data processing, management, and storage^32,33^. High-performance computing (HPC) has recently become increasingly relevant for life sciences^34,35^, especially with the availability of a user-friendly graphical user interface (GUI) through tools such as Open OnDemand^36^, which enables direct use of Jupyter notebooks and virtual desktops on HPC infrastructures^37,38^ and most importantly allows the use of elevated resources for processing of ever-growing datasets. Coupling an HPC environment with scalable data formats like OME-Zarr^39^ can help to reduce these issues while improving workflow performance. Finally, ensuring that large-scale and CLEM datasets are accessible and shareable worldwide is essential for fostering collaboration, reproducibility, and scientific progress

### Alignment

During 3D FIB/SEM image acquisition, spatial drift may occur over time due to sample charging, especially in non-conductive samples. This artefact can distort the reconstructed volume and lead to misinterpretation of object shape^40^. To prevent this, images can be aligned using “template matching” (TM)^41^, “SIFT” (scale-invariant feature transform) tools^41,42^ or by using a novel approch, such as AMST^20^ (Alignment to Median Smoothed Template), taking into consideration the local variation due to the Z FIB milling heterogeneity. Indeed, Hennies et al.^20^ demonstrate that AMST refines SIFT or TM alignments, enabling a high segmentation quality downstream. In some FIB/SEM systems, fiducial markers, often implemented as stripes or fork-shaped on top of the surface of the sample, are used to further improve dataset alignment. Thus, their presence or absence directly influences the choice of alignment strategy.

### Denoising (optional)

Denoising improves data quality while preserving information. Although optional, its usefulness depends on dataset quality and the segmentation method employed, often leading to more accurate segmentation and analysis. We selected Noise2Void2^21,43^ due to its robust denoising capabilities and its availability through both command-line tool (CLI) and GUI implementations, notably within Napari.

### Registration

CLEM datasets require precise multimodal registration, especially in 3D. A common strategy is to use landmarks or objects visible in both imaging modalities as fiducials for registration. Among available approaches^10,19^, BigWarp^18^ was selected for CLEM registration due to its precise 3D cross-correlation and its integration within BigDataViewer, a powerful tool for visualizing large 2D and 3D datasets^44^.

### Segmentation

Even with improved imaging and denoising strategies, segmentation presents additional challenges. Although it offers unprecedented insight and quantitative information in 3D volumes, it remains time-consuming and computationally demanding, and most segmentation techniques still require prior noise reduction. To address these limitations, AI-based models and neural networks, such as U-net^45^ or SAM^24^ (Segment Anything Model) have emerged as promising solutions to simplify and automate the segmentation pipeline. These models learn from manually segmented regions of interest (ROIs) and can offer accurate automated segmentation across entire datasets. Consequently, they have been increasingly applied in EM and CLEM contexts^3,11,17,46^, for instance, to segment mitochondria and other organelles using such tools through different dedicated software^19,23,26,47,48^. However, many of these steps are not user-friendly, often require programming expertise, and are not always open-souce. Napari now offers more approachable tools, such as µSAM^24^, Empanada^23^, and nnInteractive^47^, that enable automated segmentation of mitochondria and other organelles and can directly serve as input for automated CLEM registration tools, such as, CLEM-Reg^22^.

### Visualisation

Visualization is essential for communicating and interpreting results, requiring tools capable of handling 3D CLEM datasets and their segmentations in a single environment. To our knowledge, Blender is the only open-source software that enables flexible integration, visualization, and animation of complex 3D CLEM data, thanks to the Microscopy Nodes introduced by Gros et al^31^, which made it a natural choice for our workflow.

Here we introduce a modular, end-to-end solution that integrates existing and newly developed tools into a coherent pipeline for 3D CLEM processing. Our approach emphasizes flexibility by offering interoperable modules that can be combined as needed. In particular, we provide a Python-based 2D slice-alignment tool capable of FIB/SEM alignment using fiducial marks, Z-compensation, and AMST for local alignment. For datasets lacking fiducials, we also introduce an approach based on a new version of AMST (AMST2) which introduces a novel pre-alignment approach for initial coarse alignment (Hennies et al, to be published). Precise registration between LM and EM and intuitive 2D visualization are achieved using BigWarp embedded within the MoBIE^27^ plugin. For (semi-)automatic segmentation of mitochondria in an ROI, we retrained and optimised the utilisation of the open-source neural network model MitoNet. Using Empanada, it integrates smoothly into our CLEM workflow, offering a practical and accessible solution for automated segmentation. High-quality 3D rendering and animation of volumes and segmentations are performed through the Microscopy Nodes plugin in Blender^25^, completing the workflow with a powerful visualization solution. All components are open-source and we demonstrate their scalability from run efficiently from standard workstations to HPC environments. While certain steps may still require manual intervention, the ability to automate individual components on HPC infrastructure ensures scalability and efficiency. Through graphical user interfaces, comprehensive tutorials, and accessible test datasets, we empower researchers to adopt 3D CLEM workflows with confidence. By railroading the approach for user-based projects, we are streamlining and standardizing the overall process. Consequently, we provide facilities with a robust set of resources that make CLEM projects more approachable, reproducible, and manageable. Additionally, by exemplifying the use of all of the tools in high-performance environments, we demonstrate the readiness for processing of large data.

## 2. Material and methods

### 2.1. CLEM Data acquisition and collection

In this study, two different CLEM datasets from different sources were studied.

The first dataset (dataset 1 – MEFs cell with membrane contact sites (MCS)) was generated using our in-house CLEM workflow^49^, in collaboration with Wim Annaert Lab (Vrancx et al., to be published). Following earlier publications^50^, Mouse embryonic fibroblasts (MEFs) were grown on gridded glass-bottom dishes, co-expressing LAMP1-mCherry (lysosome) and GFP-Sec61β (ER labeling). Mitotracker served as a fiducial marker to align the SIM image with EM ultrastructure. Cells were fixed and imaged by 3D SIM, with an inverted Zeiss Elyra 7 (with lattice SIM module for structured illumination) microscope, equipped with two PCO.edge 4.2 CLHS sCMOS cameras in combination with a 63x oil objective (NA 1.46). The setup was controlled by ZEN black (software version (3.0), Carl Zeiss Microscopy GmbH). After SIM imaging, EM sample processing included “en bloc” staining with OTO techniques (1% Osmium & 1.5% potassium ferrocyanide, 0.2% Tannic Acid, 1% Osmium), overnight staining with 1% Uranyl acetate, and embedding in Spurr resin. After polymerization, FIB-SEM imaging was performed using a Zeiss Crossbeam 540 system with Atlas5 software on the resin blocks. Imaging was done at 1.5 kV and 600 pA using an EsB (energy-selective backscattered) detector and milling with a probe current of 700 pA). A volume of 10×20 µm with a cubic resolution of 6 nm and presenting fiducial marks was acquired following the Atlas5 procedure.

The second dataset (dataset 2 – HeLa cells) was acquired at University Medical Center Utrecht, Cell Microscopy Core using their CLEM workflow as described in Fermie et al^5,51,52^. Shortly, HeLa cells were cultured in complete DMEM and transfected with a LAMP1-GFP construct to label endo-lysosomal compartments. Cells were grown on carbon-coated gridded coverslips to enable correlative imaging. The cells were then chemically fixed and imaged using a widefield fluorescence microscope (Deltavision RT widefield microscope, GE Healthcare), a 100×/1.4 numerical aperture (NA) oil immersion objective and a Cascade II EM-CCD camera (Photometrics) with a gain value of 290 using the Acquire3D module in Softworx 6.5.2. A Z-stack was recorded for all fluorophores (LAMP1-GFP–labeled compartments) with an exposure time of 100 ms. After fixation, the samples were post-fixed and heavy-metal stained using an OTO protocol (1% osmium tetroxide and 1.5% potassium ferrocyanide, 1% thiocarbohydrazide (TCH), and 1% osmium tetroxide). The samples were subsequently stained with 2% uranyl acetate for 30 min in the dark, washed, and finally incubated with Walton’s lead aspartate (pH 5.6) for 30 min at 60 °C to further enhance contrast. The samples were then dehydrated through a graded ethanol series and infiltrated with Epon resin prior to polymerization. Finally, regions of interest were identified using the embedded grid pattern. FIB-SEM tomography was then performed using a Helios G3 UC FIB-SEM (Thermo Scientific) with a cubic voxel size of 5 nm. Serial sectioning was carried out at 30 kV with a beam current of 0.5 nA, while imaging was performed at 2 kV and 0.2 nA using the in-column backscattered electron detector. This dataset does not contain fiducial markers.

### 2.2. EM data pre-processing

To enable CLEM, a dedicated image processing workflow is required. The first step involves pre-processing of the EM data, mainly due to the inherent variability and instability of the FIB/SEM system affecting the 3D EM volume^1^. Preprocessing involves two main steps: sequential stack alignment, which ensures spatial consistency of EM images, and denoising, which enhances data quality for accurate analysis. All pre-processing steps were carried out as Slurm jobs on the Flemish supercomputer2 (Tier1 and Tier2-KUL), using GPU nodes for the denoising part. Tier1 and Tier2 KUL correspond to the regional cluster of Flanders and the university cluster associated with KU Leuven, respectively. Hence, the preprocessing steps are portable and scalable, making them interoperable and reproducible.

#### 2.2.1. 3D FIB/SEM sequential stack alignment

Different slice alignment strategies are applied depending on the presence or absence of fiducial marks in the dataset.

For datasets containing fiducial marks as generated from the Zeiss-Altas5 software, the images were processed by combining the in-house tool Taturtle and the AMST^20^ algorithm. Taturtle carries out template matching by combining the use of fiducial marks (here, stripes on top of the sample, see Figure 2) with thickness correction, ensuring a precise alignment of fiducial marks across the sample. The thickness correction consists of a per-slice z-drift correction based on recorded values during acquisition to compensate for axial shifts. This approach enables an accurate representation of the morphology of different objects within the volume along the z-axis^28^. Additionally, datasets were cropped to optimize the data treatment - this step is generally optional within the workflow. Taturtle is released as a Napari plugin with GUI and as a Nextflow^48^ module so that it can be used interactively or as a command-line tool. Subsequent to Taturtle, the AMST algorithm was applied to further improve the slice alignment by introducing non-linear transformations and thus accounting for local variations between the sample surface and depth, which are inherent to the z-milling process in 3D FIB/SEM acquisition^20^.

For datasets that do not contain fiducial marks, images were aligned using the AMST2 package (Hennies et al, to be published, https://github.com/jhennies/AMST2), which is a re-implementation of the AMST tool. In contrast to the original AMST implementation, AMST2 includes a pre-alignment step that finds translation transforms for each slice on the basis of adjacent slices as well as slices with a certain distance (here we used a distance of 8 slices). In this way, the cumulative bias along a dataset is reduced, and overall sample morphology is better preserved. In conjunction with the original AMST functionality to correct local misalignments by introducing non-linear transformations on the basis of the z-median smoothed pre-alignment dataset (i.e., the template), AMST2 yields much more accurate automatic boundary segmentation of volumetric structures^20^.

To evaluate the alignment performance, we computed displacement errors as described in Hennies et al^20^ in three local crops at both the top and bottom regions of the sample. This approach allows assessment of potential distortions along the depth of the dataset, which may result from uneven pixel-size distribution along the Z-axis^20^. For each crop, local alignment was performed, and the resulting displacements were recorded and averaged within each region. The average magnitude of these displacements was plotted and served as a quality indicator of the alignment. Further statistical analysis is presented in Figure S1 of the supporting information. All alignments were performed after Z-compensation to ensure a fair comparison.

#### 2.2.2. 3D FIB/SEM dataset denoising

Aligned datasets can be further denoised by using the CAREamics Pytorch library^53^, developed by Human technopole Italy. Specifically, we employed the Noise2Void2 (n2v2)^54^ model from CAREamics for denoising, as it corrects the artifacts introduced by the original Noise2Void approach. The denoising was run as a Python script with CAREamics packaged inside a container to promote reproducibility and interoperability on several high-performance computing clusters^54^. It should be noted that N2V2 is also available with a GUI through Napari. We used randomly selected crops over the whole dataset for the training and validation datasets to enhance computational efficiency and reproducibility.

### 2.3. 3D CLEM registration

MoBIE^27^ was used for visualization while BigWarp^18^ was employed for interactive landmark-based registration of EM and LM datasets, both integrated into BigDataViewer^55^. Prior to registration, the EM datasets were cropped to remove black borders and reduce file size, while LM channels were split and harmonized with EM in terms of pixel units and image type (8-bit). Between 10 and 20 landmarks were placed on corresponding features in both modalities, using mitochondria as fiducial marks. Affine or thin-plate spline transformations were applied depending on the dataset, with EM volumes transformed to match LM data. This choice was based on the assumption that LM represents the closest approximation to the native sample state, as EM preparation steps (e.g., dehydration and embedding) introduce sample deformation^56,57^. The transformed EM datasets were stored as a “moving image” with EM resolution and field of view, preserving origin values for accurate repositioning in the LM datasets. The registered datasets were then exported for subsequent segmentation and 3D visualization.

### 2.4. Segmentation

To exploit the CLEM dataset, segmentation of regions of interest (ROIs) was performed. Different strategies were used: (i) Automatic segmentation using Empanada-MitoNet with either the default or (ii) retrained model. In the case of retraining a model, a ground truth (GT) of the ROI was created by semi-manual segmentation using the nnInteractive plugin^47^ complemented by manual adjustment in Napari. To improve Empanada’s performance, the image contrast was inverted, as most of its training data was based on TEM images with inverted contrast^23^.

Both the datasets and GT annotations were extracted into 2D XY or XY/YZ/XZ (orthogonal) patches through an automated process implemented in a custom script, with a user-friendly interface. Both the datasets and GT annotations were extracted into 2D XY or XY/YZ/XZ (orthogonal) patches using an automated process with a custom script that features a user-friendly interface. To identify the most effective configuration, we compared different parameter combinations, such as orthogonal patches and plane usage (“run ortho plane”). Performance metrics were extracted by evaluating outputs compared to the ground truth, enabling identification of the optimal parameters (see Figure S2 in the Supporting Information).

Then, default settings were used to retrain the MitoNet model on a representative crop of the datasets and the resulting segmentation was validated by extracting quantitative metrics through comparison with a ground truth. Subsequently, the retrained model was used to segment the full datasets, which were further refined using Napari tools (bucket, label 0 and “edit dim” of 3) to remove inaccurate labels.

### 2.5. 3D visualisation and animation using Microscopy Nodes and Blender

CLEM datasets were finally visualised and animated using the version 4.5.1 of Blender© with the Microscopy Nodes^25^ and expanded to the use of CLEM data. LM datasets were visualised by volumetric and emissive renders, while the registered EM dataset is rendered as light-scattering volumes. The segmentation of the object of interest was visualised by a 3D mesh and displayed as a spatial mask. To balance visibility between emissive LM and scattering EM, transparent or gray backgrounds were applied. Precise 3D positioning was achieved through axis translation, ensuring accurate overlay of EM and LM datasets and correct placement of mesh annotations. Therefore, saving ROI coordinates and origin values of registered images is essential for reproducibility. Visualisation and animation were obtained by following the channel tutorial^25^.

### 2.6. Data management and availability

During the project, the data was transferred using Globus and backed up on the active data management platform ManGO (https://rdm-docs.icts.kuleuven.be/mango/index.html), developed by KU Leuven. We have deposited the raw and the processed data on the Bioimage archive available at [S-BIAD2993].

The Taturtle Napari plugin, Python scripts, and the associated Nextflow module are available under a BSD-3-Clause license at https://github.com/vib-bic-projects/2025_10_Helene.git.

## 3. RESULTS

To develop a turnkey analysis solution for 3D CLEM datasets, we have been evaluating the entire workflow from processing to visualization. By combining existing tools with new ones, we built an open-source end-to-end solution for 3D CLEM datasets. Consequently, the workflow incorporates evaluation of alignment strategies with or without fiducial marks and the optimization of automated segmentation using Empanada-MitoNet. The final step offers high-quality 3D visualization and animation through Microscopy Nodes. Figure 1 illustrates the flow and modularity of the solutions. In the following, we describe and evaluate the individual parts of the workflow and how they can be combined.

**Figure 1:**
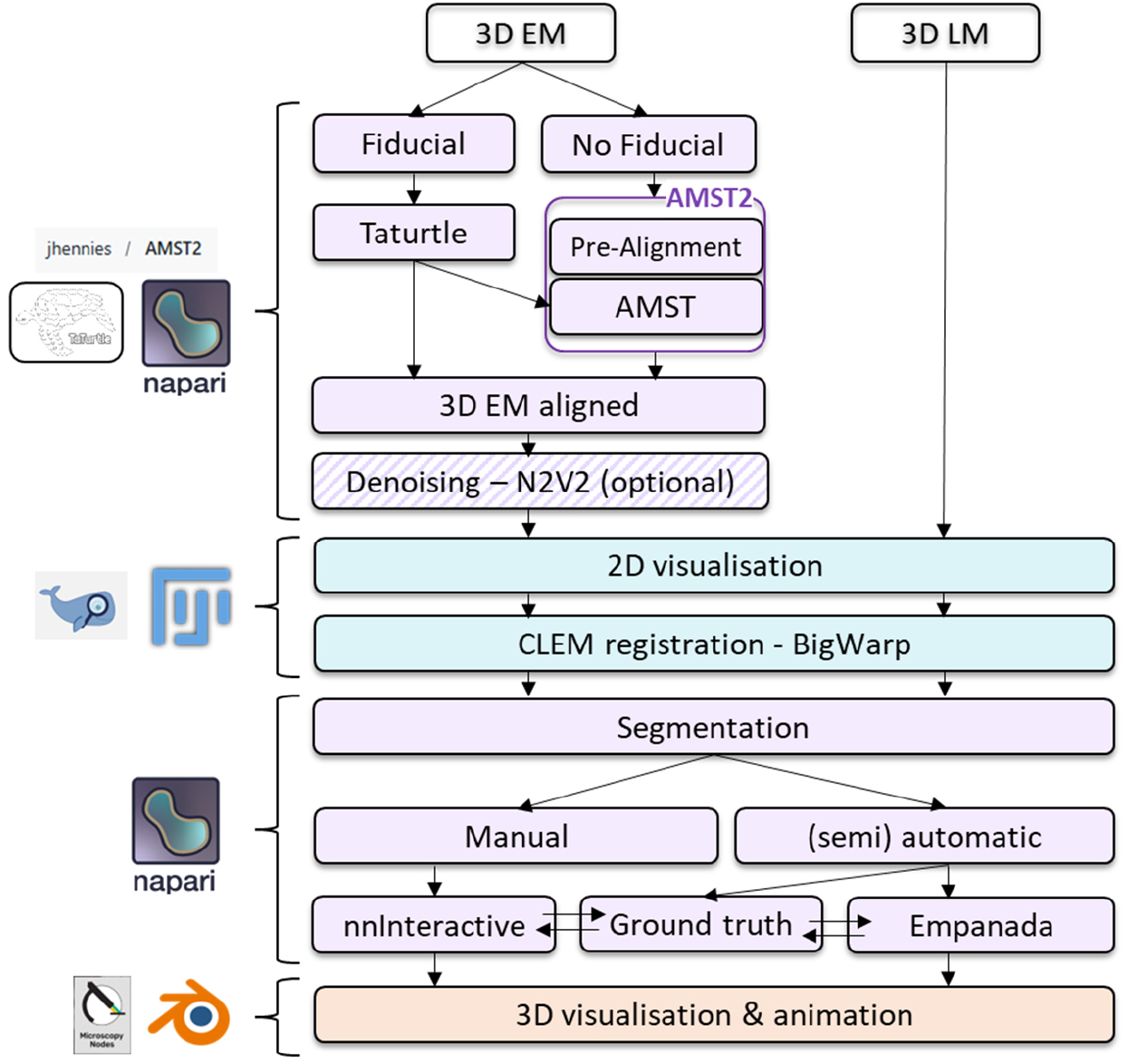
Open-source image processing workflow to process CLEM datasets. From top to bottom: preprocessing (purple), segmentation (green), and 3D visualisation (orange) steps supported by Fiji, Napari, and Blender, respectively.

### 3.1. Evaluation of alignment strategies for 3D FIB/SEM datasets with fiducial marks

To enhance visualization quality and segmentation accuracy, we compared the performance of several alignment strategies using the 3D FIB/SEM dataset 1 (Figure 2). This dataset includes fiducial marks in the form of black stripes introduced during platinum deposition in the milling direction (Z-axis of the later dataset), which are clearly visible on the top layer in the XY plane (Figure 2a). Hence, the marks can serve as structural references for aligning image slices throughout the volume.

**Figure 2:**
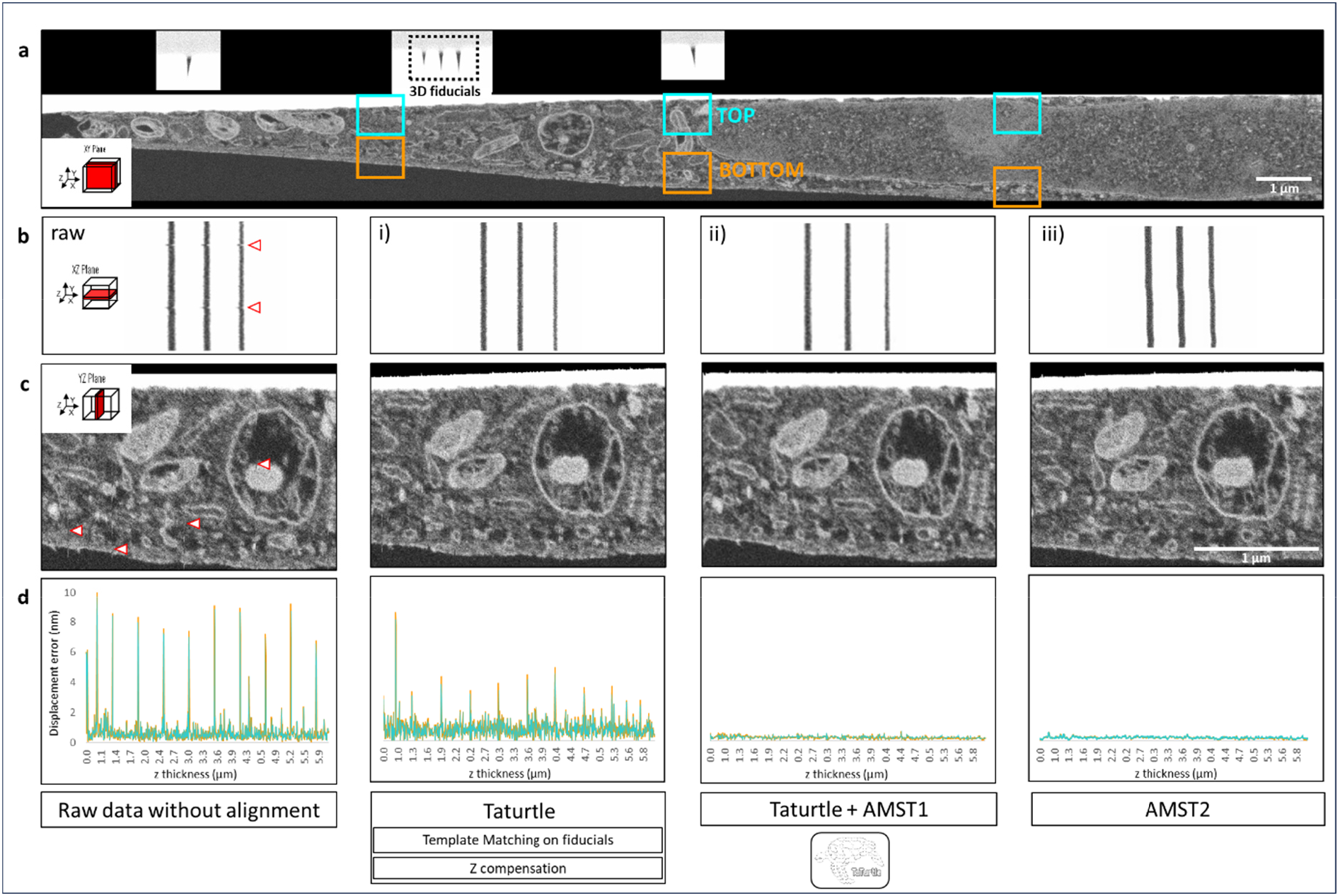
Comparison of different strategies for alignment of dataset 1 - a 3D FIB/SEM dataset containing fiducials marks: a) XY plane of the raw dataset, where black fiducial marks are visible on top of the platinum deposition; b) XZ plane, top view of the fiducial marks and c) XY plane – display the dataset before (i) and after alignment using three approaches: Taturtle (an in-house plugin combining Z-compensation and template matching on fiducials) (ii), Taturtle combined with AMST (Alignment to Median Smoothed Template) (iii), and, AMST2 (iv). For each alignment strategy, b and c highlight the improvements in stack alignment (with artifacts indicated by red arrows), and d presents the average displacement errors measured in the top (cyan rectangles) and bottom region (orange rectangles) of the dataset. All plots share the same y-axis. Note that most of the periodic peaks of high alignment error present in the raw data and Taturtle result coincide with auto-focus and auto-stigmation operations during image acquisition.

We compared three alignment tools: (i) Taturtle, our in-house plugin combining Z-compensation with template matching on fiducials; (ii) Taturtle combined with AMST; and (iii) AMST2s pre-alignment combined with AMST. For readability the tools will in the following be referred to as (i) Taturtle, (ii) Taturtle-AMST and (iii) AMST2.

All strategies substantially improve the fiducial mark alignment by eliminating small shifts observed in the view of the raw data, indicating enhanced consistency along the Z-axis (Figure 2b, red arrows). Respective improvements in alignment are also evident in the XY view, by observing continuous organelle membranes (Figure 2c). Visually, AMST2 and Taturtle-AMST yield an improved continuous stack alignment, suggesting effective correction of geometric distortions across slices (Figure 2b and c). Additionally, we computed displacement errors as described in Hennies et al^20^. In line with previous results, Taturtle-AMST and AMST2 alignment achieved similar performance and demonstrated clear improvements over raw data and Taturtle (from ≈0.75 nm to ≈0.25nm of displacement error in Z, see statistical result in supporting information). In contrast, no noticeable difference in displacement error was observed between the analysed regions (top and bottom; see supporting information). This likely reflects the flat morphology of the cell (only a few micrometres thick), which minimizes distortion caused by deep FIB milling.

To further assess the robustness of AMST2, we applied the method to a version of the same dataset without fiducial marks. Although the resulting alignment appeared visually acceptable, closer inspection revealed subtle deformation artifacts: the cell volume displayed a slightly convex surface, and the cell base was no longer flat against the coverslip, which we assume to be flat under normal conditions (see Figure S3 in supporting information). However, it remains unclear whether such an artefact systematically affects all volume EM datasets processed without fiducials, as this likely depends on sample geometry and acquisition conditions. Overall, while AMST2 performs well in the absence of fiducial marks, however, our findings suggest that the presence of fiducial marks can improve alignment accuracy.

In summary, both Taturtle-AMST and AMST2 offer reliable alignments and produce similar visual outcomes. AMST2 has the advantage of delivering satisfying results even when fiducial marks are absent. However, performance may vary depending on factors such as image quality, modality, and contrast. Additionally, in contrast to Taturtle, this tool is not available with a GUI yet. Therefore, the choice of alignment tool and strategy should be adapted to the specific characteristics and heterogeneity of each dataset.

### 3.2. Alignment and denoising enhance automated mitochondrial segmentation

For dataset 2, mitochondria, the objects of interest, were segmented across the entire dataset using the automatic segmentation approach with Empanada-MitoNet^23^.To evaluate the impact of alignment and denoising on automated segmentation, we tested their effects on this dataset, which is more representative of typical FIB/SEM acquisitions. In opposition to dataset 1, dataset 2 displayed/showed greater structural variability and lacked fiducial marks, making it a more challenging and realistic scenario for testing segmentation robustness. Applying pre-processing under these conditions allowed us to assess the impact of alignment and denoising on segmentation performance within the workflow, suggesting that these steps could be implemented routinely. We performed this analysis using AMST2 stack alignment and N2V2 denoising to segment mitochondria with the *MitoNet* model^23^ (Figure 3).

**Figure 3:**
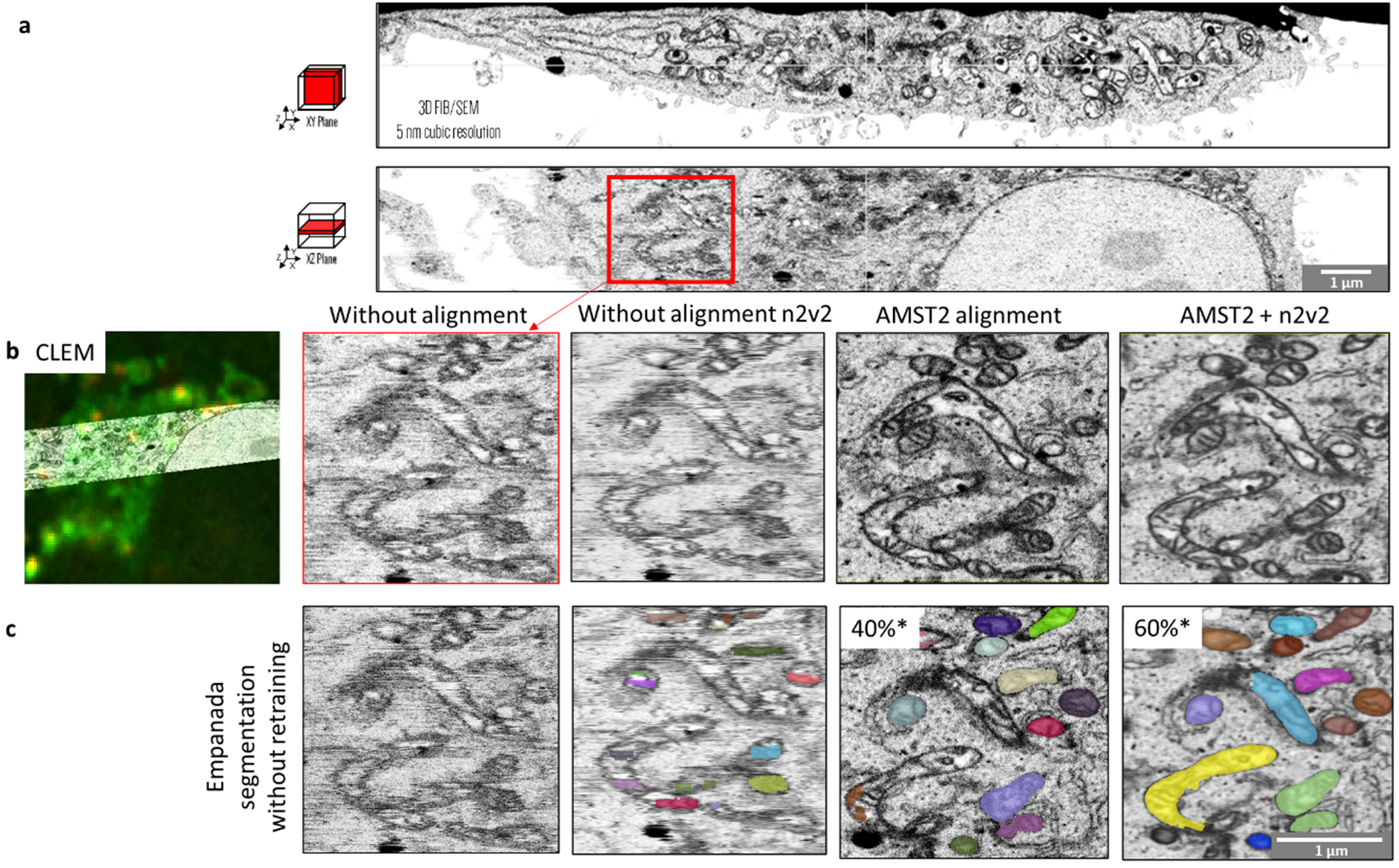
Impact of AMST2 alignment and N2V2 denoising on automated mitochondrial segmentation in a CLEM workflow. a) The native 3D FIB/SEM dataset in XY and YZ planes. The enlarged region (red rectangle) highlights drift and milling-related artifacts along the Z-axis. b) the same region in the CLEM context and displays, from left to right, the results after applying N2V2 denoising, AMST2 alignment, and the combination of both methods. c) the corresponding mitochondrial segmentations generated with Empanada-MitoNet (default parameters, no additional retraining). Quantitative metrics comparing segmentation outputs to ground truth are also provided (*).

The raw dataset in both XY- and XZ-planes revealed the structural distortions commonly observed in unprocessed volumes (Figures 3a). Pre-processing using AMST2 alignment and/or N2V2^54^ denoising resulted in a reduction of noise while fine structural details were preserved (Figure 3b). These processed datasets served as input for Empanada-MitoNet with default settings and without model retraining. The combined application of AMST2 alignment and N2V2 denoising resulted in a reduction of noise while fine structural details were preserved (Figure 3c). Segmentation accuracy improved from approximately 40% with alignment alone to nearly 60% when combined with denoising, confirming the critical role of preprocessing in optimizing automated segmentation performance. Note that segmentation parameters had to be adapted due to the variability of EM datasets.

Despite the absence of fiducials in this dataset, the applied strategy provided strong results, suggesting that such an approach may be extended to other challenging FIB/SEM datasets and modalities. These findings highlight the potential of alignment and denoising workflows to improve segmentation outcomes and provide more perspectives in correlative imaging contexts.

### 3.3. Optimised model retraining enhances automated mitochondrial segmentation while minimizing pre-processing requirement

Given previous results, we next investigated model retraining as a complementary strategy to further improve segmentation performance. We retrained Empanada-MitoNet prior to its application on the full dataset. This approach significantly improved segmentation accuracy compared to the untrained model (Figure 4).

**Figure 4:**
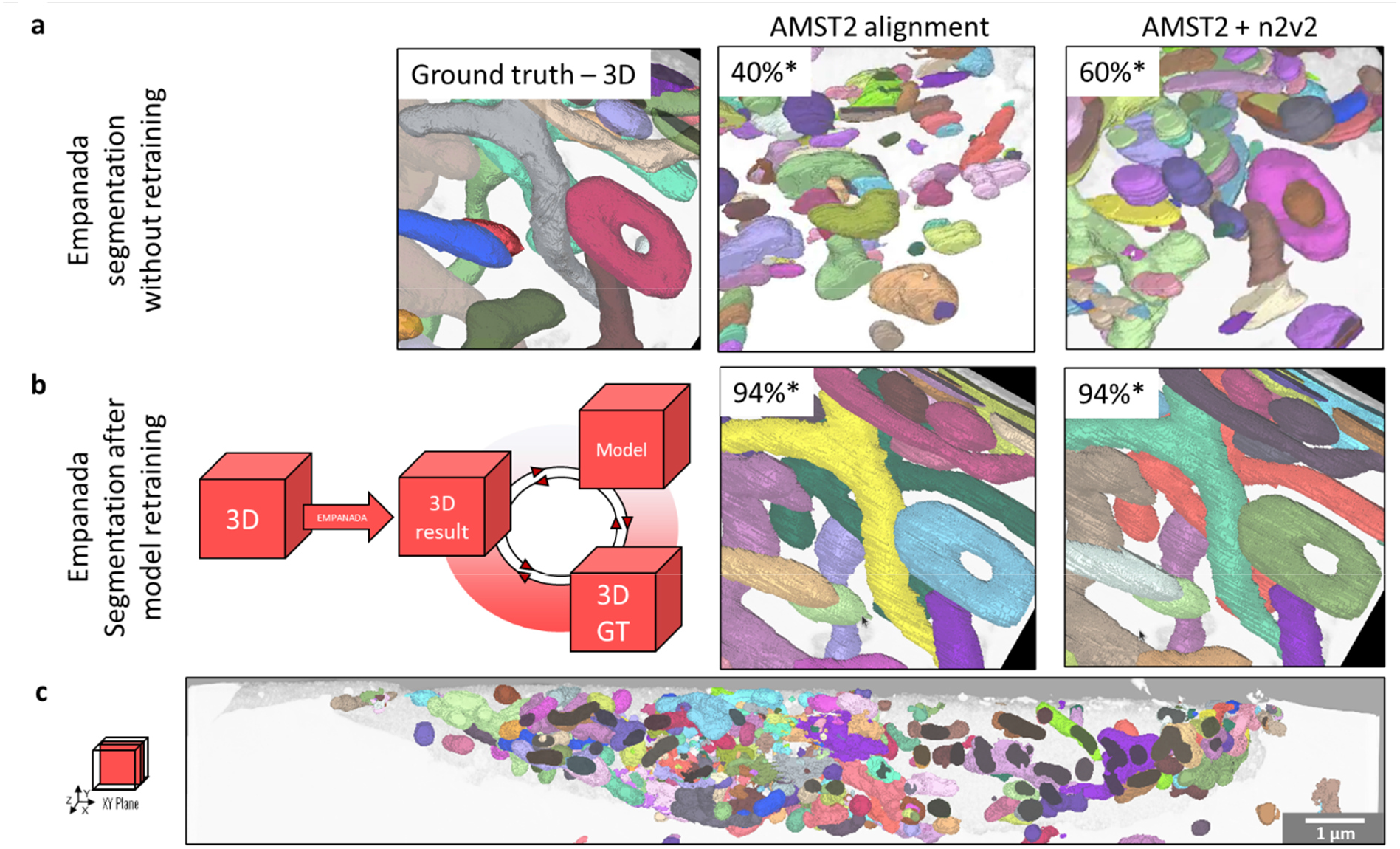
Model retraining improvement on mitochondrial segmentation using Empanada. a) 3D visualisation of segmentation generated without model retraining from the same ROI of the dataset 2 (see Figure 3), aligned with AMST2, and optionally denoised using N2V2. Alongside, its ground truth (GT) annotations were created using the nnInteractive plugin in Napari. b) Segmentation results after model retraining, illustrating the strategy proposed in this study: the GT annotations were used to retrain the Empanada model prior to its application on the full dataset. Quantitative metrics comparing segmentation outputs to ground truth are also provided (*). c) Automatic segmentation of the full dataset after model retraining.

As shown previously, segmentation without model retraining reveals fragmented segmentation in comparison to the ground truth, even after alignment with AMST2 and optionally N2V2 denoising (Figure 4a). After retraining of the model, 3D visualisation of the segmentation revealed a substantial improvement (Figure 4b). This was confirmed by the segmentation accuracy, increasing from approximately 40% to nearly 94% after retraining. This improvement was consistent for both aligned datasets with and without denoising, demonstrating the robustness of the retrained model. The results correspond to the best-performing configuration based on 2D patches without orthogonal-plane runs (see Experimental section). Finally, the retrained model successfully segmented the mitochondria of the full dataset (Figure 4c).

This work demonstrates that model retraining not only enhances segmentation performance and accessibility for non-programming users but also achieves accuracy levels where denoising has no additional effect, underscoring its critical role in high-precision segmentation.

### 3.4. BigWarp and BigDataViewver provide a rapid and robust solution for CLEM volume registration and 2d visualisation

To achieve an accurate correlation between the LM and EM datasets, the datasets were registered with the BigWarp^18^ plugin embedded within BigDataViewer^55^. Here, the mitotracker channel of the LM stack and the morphology of the mitochondria in EM were used as fiducials and landmarks (Figure 5). The resulting transformation and landmark were visualized using the MoBIE-plugin. This workflow demonstrates that combining BigWarp with MoBIE provides an efficient and reproducible solution for rapid and accurate registration and visualization of CLEM datasets.

**Figure 5:**
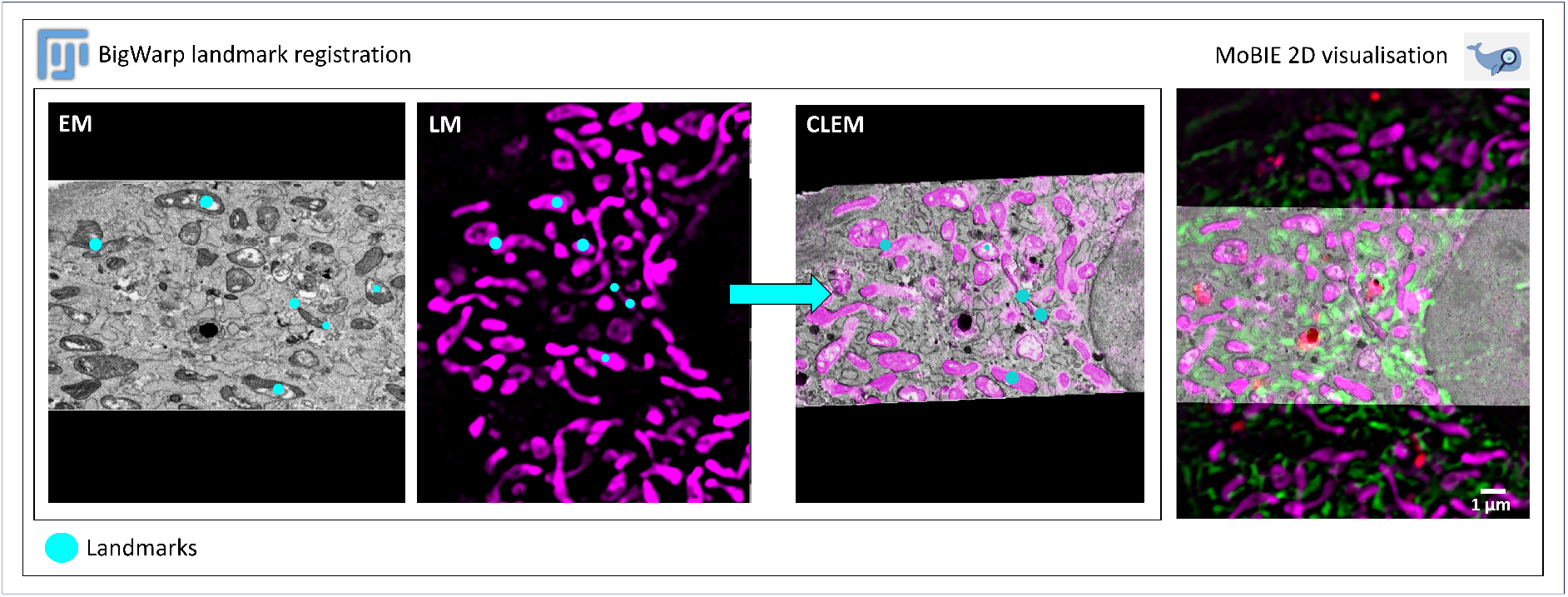
BigWarp landmark-based registration and 2D visualization of a 3D CLEM dataset. The left panel illustrates the registration workflow, where landmarks (cyan dots) were placed on corresponding details in LM (mitotracker signal in magenta) and EM (mitochondrial morphology) datasets using mitochondria as fiducial marks within BigWarp embedded in BigDataViewer. The right panel shows the registered dataset visualized as 2D orthogonal planes in BigDataViewer.

### 3.5. Microscopy Nodes provide high-quality animation and 3D visualization of CLEM dataset and segmentation in a single environment

Following all previous steps, we achieve full 3D visualisation of the CLEM dataset 1. Using Microscopy Nodes (MN)^25^ embedded in Blender, we generated both images (Figure 6) and an animated video (video S1 in support information).

**Figure 6:**
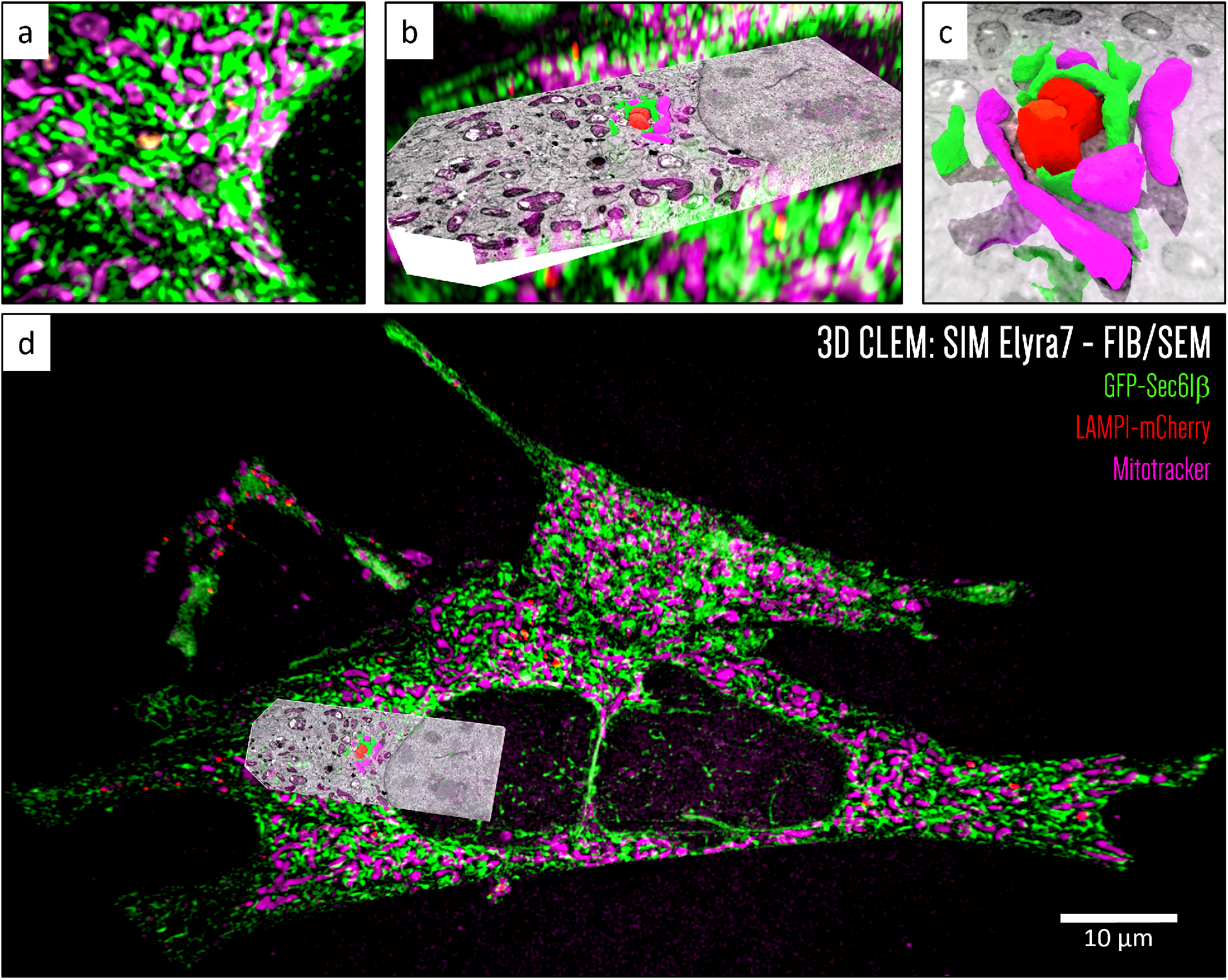
3D CLEM rendering of a large dataset using Microscopy nodes. A) Zoomed-in view of a 3D SIM dataset of a MEF cell, showing the ER (green), late endosomes/lysosomes (red), and mitochondria (magenta). b) The same region imaged by a FIB/SEM and registered with LM, shown as a 3D reconstruction rendered at high isotropic resolution (6×6×6nm), revealing the finest structural details of the volume. c) 3D segmentation and rendering of a selected region, highlighting potential membrane contact sites (MCS), visualized in three dimensions. The same color code as the LM channels was applied for consistency. This segmentation is also shown in Figure 6b. d) Overview of the sample, illustrating the integration and spatial registration of the different imaging modalities within the CLEM workflow.

Microscopy Nodes (MN) enables the integration, visualization, and animation of complex 3D CLEM datasets within a single environment. It also supports large datasets, which are common in CLEM workflows due to their multimodal nature. Microscopy nodes support both TIFF and OME-Zarr formats^39^. For dataset 1, only a small area required segmentation. Based on previous studies^49,50^, a region of interest (ROI) containing potential membrane contact sites (MCS) was selected. Mitochondria, ER, and lysosomes within the ROI were segmented using the semi-automated method.

The 3D rendering of the dataset 1 includes: a 2.5 GB 3D SIM dataset of a MEF cell imaged in three fluorescent channels (see Experimental Section) (Figure 6a); a 17 GB EM volume registered using mitochondrial landmarks visible in both LM and EM, acquired via 3D FIB/SEM at 6 nm isotropic resolution (Figure 6b); and a 675 MB segmentation of a region of interest rendered as a 3D mesh (Figure6c). In the visualization environment, LM data are displayed with each voxel emitting light proportionally to its recorded fluorescence intensity, reproducing the original acquisition conditions^25,30^, whereas the EM volume is represented as a dense block to reproduce the appearance of SEM images acquired on the FIB trench surface. The segmented ROI is incorporated directly into the 3D scene as a mesh, enabling smooth co-visualization with both LM and EM volumes (Figure 6d).

This workflow demonstrates the robustness of Microscopy Nodes as an open-source, powerful 3D visualization and rendering tool, compatible with big dataset management and HPC usage, effectively concluding our CLEM image-processing workflow and enabling high-quality animations (see Supporting Information).

## 4. Discussion

CLEM workflows are powerful approaches and, in some cases, the only suitable way to address specific scientific questions. Despite growing demand, processing and analysing 3D CLEM datasets remains challenging due to their complexity, multimodal nature, and extensive post-processing requirements, which make them difficult to implement within microscopy facilities as a service. The lack of standardized workflows and the fragmented landscape of image-processing tools, particularly in critical steps such as FIB/SEM slice alignment and automated segmentation, hinder such development. In addition, it requires managing a vast amount of data. Thus, an open-source image-processing workflow combined with the usage of High-performance computing (HPC) and Jupyter notebooks offer a promising approach towards broader accessibility and scalability. Consequently, similar strategies may lead to broader adoption, not only in the CLEM field but also in other types of correlative imaging and across multiple disciplines.

We present a complete end-to-end open-source workflow for processing, visualizing and animating 3D CLEM datasets, along with two new tools: (i) a slice-alignment tool for FIB/SEM data with and without fiducials marks, that integrates Z-compensation, and AMST; and (ii) a methodology for optimizing automatic segmentation of mitochondria in Empanada-MitoNet through parameter choice and metrics evaluation. And (iii) demonstrate the robustness of our workflow by showing successful 3D CLEM high-quality visualisation and animation through Microscopy Nodes. Moreover, we demonstrate the scalability of our solution by using our solutions on HPC platforms.

To ensure that our image processing workflow is accessible and generalizable to the broader CLEM community, we included datasets with and without fiducial marks (see Figures 1 and 2). Using Taturtle with AMST or AMST2 as alignment tools, we improve dataset geometric consistency across slices, even in the absence of fiducial marks (see Figures 1 and 2). However, while AMST2 remains effective, our results suggest that alignment performance could benefit from the use of fiducial marks. Overall, this ensures that our tools are both accessible and generalizable for the broader EM and CLEM communities. The combination of good alignment with Noise2Void2 denoising improves pre-processing quality, highlighting the need for accurate preprocessing for reliable visualisation and further segmentation. When model retraining is unavailable, we demonstrate that alignment and denoising are essential preprocessing steps that improve the accuracy of mitochondrial segmentation by more than 60% on 3D FIB/SEM datasets. However, it should be noted that the efficiency of pre-processing steps remains influenced by contrast, resolution, noise, and sample preparation.

In addition, our results demonstrate that model retraining is a critical step for obtaining high-quality automated mitochondrial segmentation. Segmentation accuracy improved from approximately 60% without retraining to nearly 94% after retraining, even on datasets without denoising, underscoring the robustness of the retrained network. Despite this, segmentation parameters still required adjustment due to the inherent variability of EM datasets. Because Empanada^23^ offers a wide range of parameters, it becomes unrealistic to manually test or rationally select the best configuration for each dataset. Hence, integrating a parameter-optimization strategy would, therefore, be a valuable addition to any robust segmentation pipeline across diverse EM datasets and modalities.

By registering EM to LM, we emphasize that LM was chosen as the reference modality, assuming that EM sample preparation includes structural deformation, making LM closer to the sample’s native state. By using fiducial landmarks placement and affine transforms through the BigWarp^18^ plugin, we were able to display datasets across modalities. Furthermore, we also demonstrate fast and efficient 2D visualization of CLEM datasets using BigDataViewer^44^, followed by high-quality 3D rendering and animation with Microscopy Nodes. Segmentation outputs can be registered and visualized directly within the CLEM volume, illustrating the strength and flexibility of this visualization approach. While other open-source software, such as Napari^58^, MIB^48^, or Icy^59^, also support 3D rendering and could be integrated into our workflow, Microscopy Nodes^31^ integrated into Blender was selected for its ability to display, register, and animate multiple modalities and segmentations within a single highly customizable environment. This makes it a particularly powerful option for advanced CLEM visualization.

Here, we decided to reuse and integrate mostly existing software tools in scientific workflows to enhance efficiency, reliability, and reproducibility. Established tools benefit from years of community development and optimization, allowing researchers to focus on scientific questions rather than computational challenges. Additionally, this approach improves interoperability, enabling easier sharing and reproduction of datasets and results, and avoids duplication and fragmentation of solutions, consequently fostering collaboration and transparency. Moreover, utilizing mature software ensures robustness through active user communities and regular updates, a level of support rarely matched by custom solutions. Finally, the design and flexibility of our workflow ensures scalability for large datasets (though HPC compatibility) and adaptability across research groups. Altogether, these elements position our workflow as a comprehensive and accessible solution for CLEM image processing, capable of supporting diverse biological applications and, we believe, fostering broader adoption within the community. From the perspective of expanding such a workflow remains challenging. Future improvement could include integration with live-cell imaging, which could be directly visualized and animated within Microscopy Nodes^31^. Additionally, further testing across a wider range of imaging modalities, resolutions, and biological contexts will be essential to consolidate the workflow’s general applicability.

## Supporting information

Video link video S1

## 5. Acknowledgments

Computing resources and services used in this work by S.M., B.P., H.R., and T.W. were provided by the VSC (Flemish Supercomputer Center), funded by the Research Foundation - Flanders (FWO) and the Flemish Government via a grant to Flanders BioImaging. S. M. and H.R. were supported by a grant from the FWO (FWOI000123N). In addition, S.M. was supported by grants from FWO (FWOI001322N) and KULeuven; KA/24/041. The authors gratefully acknowledge the VIB BioImaging Core Leuven and Ghent, P.Hernández-Varas and A.Kremer for their support and assistance, K. Vints for her advice regarding sample preparation, R. Sannerud for her valuable advice regarding cell management, imaging, and the study context, and Prof Wim Annaert (FWO Heavy Infrastructure grant I001322N). We acknowledge the access and services provided by the Imaging Centre at the European Molecular Biology Laboratory (EMBL IC), generously supported by the Boehringer Ingelheim Foundation. The authors thank M.Croft and J.Deschamps (Jug lab, Human Technopole Italy), for answering questions and providing support about CAREamics. We thank A.Zakieva (EMBL Heidelberg) for helpful discussion regarding data management and BioImage Archive deposition.

## Supporting information

### 3D visualisation and animation of registered large clem dataset using microscopy nodes

Video S1: 3D visualisation and video animation of correlative 3D imaging of a MEF cell combining SIM and FIB/SEM, showing the ER (green), late endosomes/lysosomes (red), and mitochondria (magenta), with high-resolution vEM reconstruction (6 × 6 × 6 nm) (grey) correlated to LM via the mitotracker signal. In addition, a 3D segmentation is highlighting potential membrane contact sites (MCS) using the same color code as the LM channels.

### Statistical analysis of the displacement error

**Figure S1:**
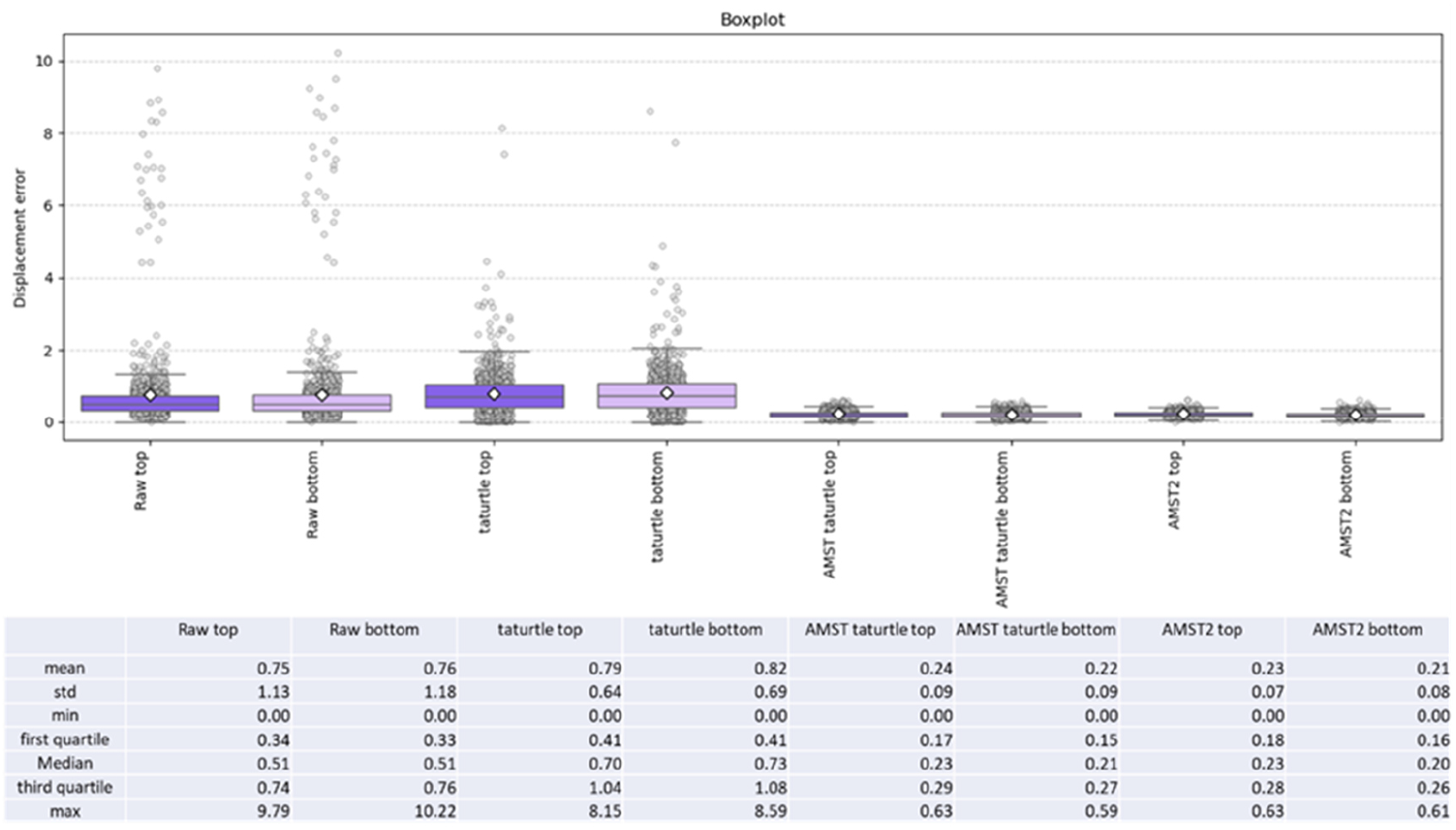
Statistical analysis of the mean displacement error for each condition and region (top and bottom). Both Taturtle and AMST2 reduce the gap between the mean (white hexagon) and the median (central line of the boxplot) and lower the number of extreme outliers (points falling outside the whiskers), demonstrating the impact of alignment strategies on raw data. By comparing the top (dark purple boxes) and bottom (light purple boxes) region, no obvious difference in displacement error was observed.

### Evaluation of segmentation accuracy using metrics extraction

**Figure S2:**
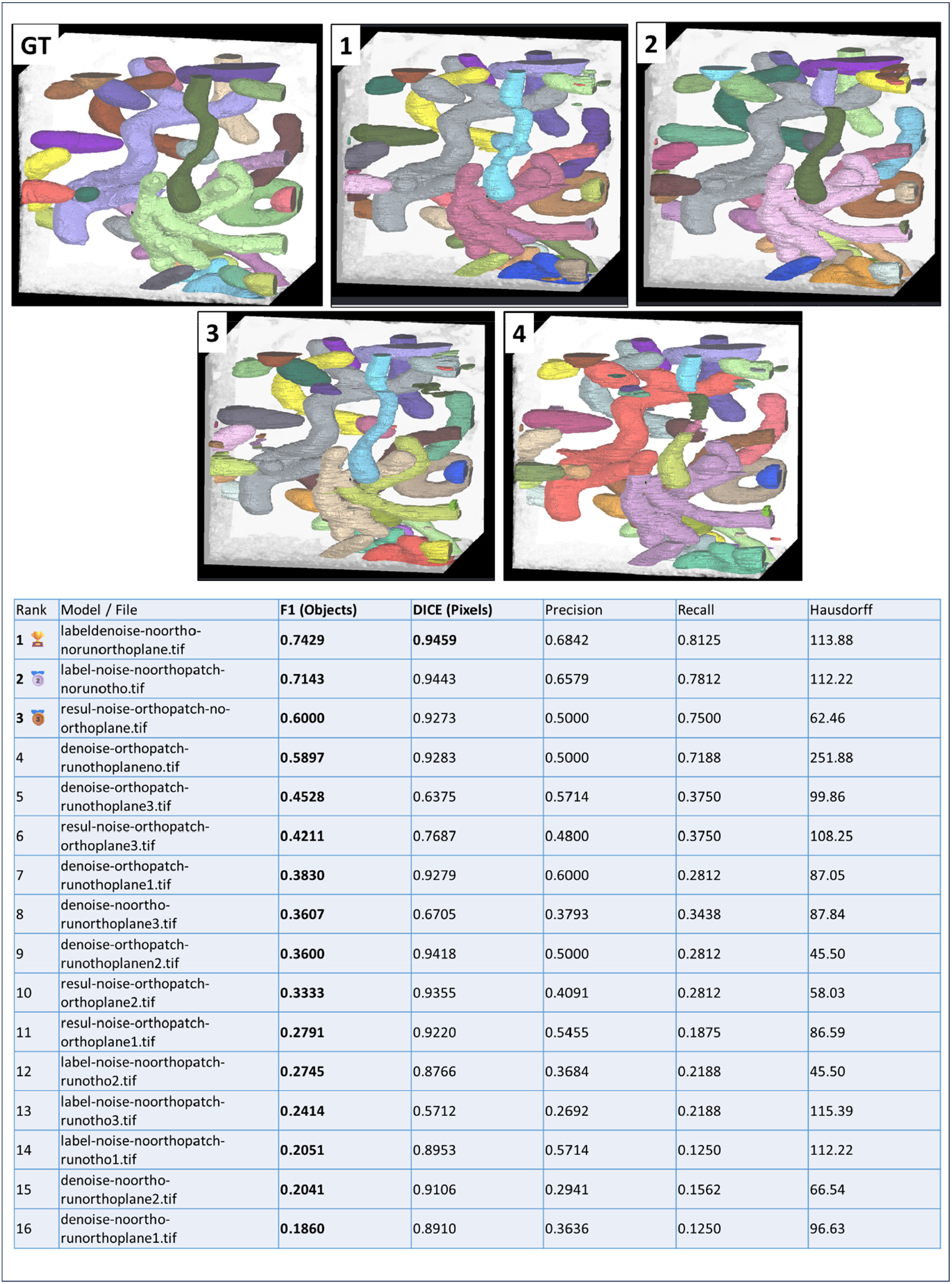
3D visualisation obtained from the 4 first best segmentation presenting the best metrics score. The ground truth of the same area is also presented as the reference of comparison.

To identify the most effective segmentation parameters configuration, we compared different parameter combinations, such as orthogonal patches and plane usage (“run ortho plane”). Performance metrics were extracted from comparison between segmentation outputs and ground truth of a representative cropped area of the dataset (see Figure S1). The DICE metrics was chosen here to represent the segmentation as it is a statistical measure used to evaluate the similarity between two sets, commonly applied in image segmentation.

### Evaluation AMST2 aligment in presence or absence of fiducials marks

**Figure S3:**
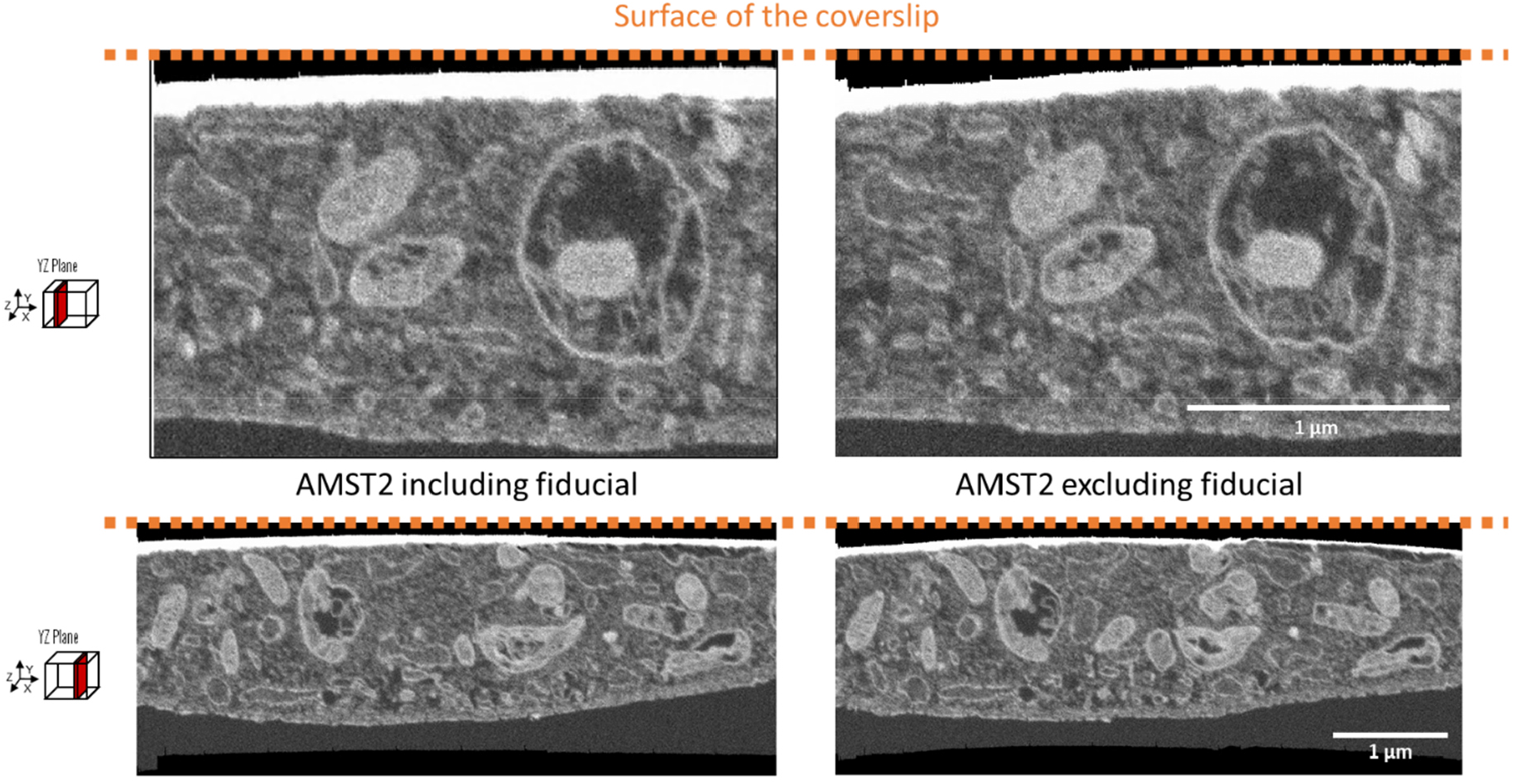
Comparison of AMST2 alignment output on a 3D FIB/SEM dataset including or not fiducials marks. With fiducial, the cell surface is flat against the coverslip surface (orange dots line), which we assure to be flat here (left panel). However, if fiducials are cropped out of the dataset, AMST2 alignment output presents a slightly convex surface (right panel).

## References

(1) Collinson, L. M.; Carroll, E. C.; Hoogenboom, J. P. Correlating 3D Light to 3D Electron Microscopy for Systems Biology. Curr. Opin. Biomed. Eng. 2017, 3, 49–55. 10.1016/j.cobme.2017.10.006.

(2) Kremer, A.; Van Hamme, E.; Bonnardel, J.; Borghgraef, P.; Guérin, C. j.; Guilliams, M.; Lippens, S. A Workflow for 3D-CLEM Investigating Liver Tissue. J. Microsc. 2021, 281 (3), 231–242. 10.1111/jmi.12967.

(3) Ohta, K.; Hirashima, S.; Miyazono, Y.; Togo, A.; Nakamura, K. Correlation of Organelle Dynamics between Light Microscopic Live Imaging and Electron Microscopic 3D Architecture Using FIB-SEM. Microscopy 2021, 70 (2), 161–170. 10.1093/jmicro/dfaa071.

(4) van Rijnsoever, C.; Oorschot, V.; Klumperman, J. Correlative Light-Electron Microscopy (CLEM) Combining Live-Cell Imaging and Immunolabeling of Ultrathin Cryosections. Nat. Methods 2008, 5 (11), 973–980. 10.1038/nmeth.1263.

(5) Fermie, J.; Liv, N.; ten Brink, C.; van Donselaar, E. G.; Müller, W. H.; Schieber, N. L.; Schwab, Y.; Gerritsen, H. C.; Klumperman, J. Single Organelle Dynamics Linked to 3D Structure by Correlative Live-Cell Imaging and 3D Electron Microscopy. Traffic 2018, 19 (5), 354–369. 10.1111/tra.12557.

(6) Johnson, E.; Seiradake, E.; Jones, E. Y.; Davis, I.; Grünewald, K.; Kaufmann, R. Correlative In-Resin Super-Resolution and Electron Microscopy Using Standard Fluorescent Proteins. Sci. Rep. 2015, 5 (1), 9583. 10.1038/srep09583.

(7) Sexton, D. L.; Burgold, S.; Schertel, A.; Tocheva, E. I. Super-Resolution Confocal Cryo-CLEM with Cryo-FIB Milling for in Situ Imaging of Deinococcus Radiodurans. Curr. Res. Struct. Biol. 2022, 4, 1–9. 10.1016/j.crstbi.2021.12.001.

(8) Hoffman, D. P.; Shtengel, G.; Xu, C. S.; Campbell, K. R.; Freeman, M.; Wang, L.; Milkie, D. E.; Pasolli, H. A.; Iyer, N.; Bogovic, J. A.; Stabley, D. R.; Shirinifard, A.; Pang, S.; Peale, D.; Schaefer, K.; Pomp, W.; Chang, C.-L.; Lippincott-Schwartz, J.; Kirchhausen, T.; Solecki, D. J.; Betzig, E.; Hess, H. F. Correlative Three-Dimensional Super-Resolution and Block-Face Electron Microscopy of Whole Vitreously Frozen Cells. Science 2020, 367 (6475), eaaz5357. 10.1126/science.aaz5357.

(9) Hegermann, J.; Wrede, C.; Fassbender, S.; Schliep, R.; Ochs, M.; Knudsen, L.; Mühlfeld, C. Volume-CLEM: A Method for Correlative Light and Electron Microscopy in Three Dimensions. Am. J. Physiol.-Lung Cell. Mol. Physiol. 2019, 317 (6), L778–L784. 10.1152/ajplung.00333.2019.

(10) Weiner, A. Step-by-Step Guide to Post-Acquisition Correlation of Confocal and FIB/SEM Volumes Using Amira Software. Methods Cell Biol. 2021, 162, 333–351. 10.1016/bs.mcb.2020.09.006.

(11) Mocaer, K.; Mizzon, G.; Gunkel, M.; Halavatyi, A.; Steyer, A.; Oorschot, V.; Schorb, M.; Le Kieffre, C.; Yee, D. P.; Chevalier, F.; Gallet, B.; Decelle, J.; Schwab, Y.; Ronchi, P. Targeted Volume Correlative Light and Electron Microscopy of an Environmental Marine Microorganism. J. Cell Sci. 2023, 136 (15), jcs261355. 10.1242/jcs.261355.

(12) Radulović, S.; Sunkara, S.; Rachel, R.; Leitinger, G. Three-Dimensional SEM, TEM, and STEM for Analysis of Large-Scale Biological Systems. Histochem. Cell Biol. 2022, 158 (3), 203–211. 10.1007/s00418-022-02117-w.

(13) Luckner, M.; Wanner, G. Precise and Economic FIB/SEM for CLEM: With 2 Nm Voxels through Mitosis. Histochem. Cell Biol. 2018, 150 (2), 149–170. 10.1007/s00418-018-1681-x.

(14) Correlative Imaging of Collagen Fibers and Fibroblasts Using CLEM Optimized for Picrosirius Red Staining and FIB/SEM Tomography. J. Electron Microsc. (Tokyo) 2020, 69 (5), 324–329. 10.1093/JMICRO/DFAA024.

(15) Marion, J.; Le Bars, R.; Satiat-Jeunemaitre, B.; Boulogne, C. Optimizing CLEM Protocols for Plants Cells: GMA Embedding and Cryosections as Alternatives for Preservation of GFP Fluorescence in Arabidopsis Roots. J. Struct. Biol. 2017, 198 (3), 196–202. 10.1016/j.jsb.2017.03.008.

(16) Booth, D. G.; Beckett, A. J.; Molina, O.; Samejima, I.; Masumoto, H.; Kouprina, N.; Larionov, V.; Prior, I. A.; Earnshaw, W. C. 3D-CLEM Reveals That a Major Portion of Mitotic Chromosomes Is Not Chromatin. Mol. Cell 2016, 64 (4), 790–802. 10.1016/j.molcel.2016.10.009.

(17) Müller, A.; Schmidt, D.; Albrecht, J. P.; Rieckert, L.; Otto, M.; Galicia Garcia, L. E.; Fabig, G.; Solimena, M.; Weigert, M. Modular Segmentation, Spatial Analysis and Visualization of Volume Electron Microscopy Datasets. Nat. Protoc. 2024, 1–31. 10.1038/s41596-024-00957-5.

(18) Bogovic, J. A.; Hanslovsky, P.; Wong, A.; Saalfeld, S. Robust Registration of Calcium Images by Learned Contrast Synthesis. In 2016 IEEE 13th International Symposium on Biomedical Imaging (ISBI); 2016; pp 1123–1126. 10.1109/ISBI.2016.7493463.

(19) Paul-Gilloteaux, P.; Heiligenstein, X.; Belle, M.; Domart, M.-C.; Larijani, B.; Collinson, L.; Raposo, G.; Salamero, J. EC-CLEM: Flexible Multidimensional Registration Software for Correlative Microscopies. Nat. Methods 2017, 14 (2), 102–103. 10.1038/nmeth.4170.

(20) Hennies, J.; Lleti, J. M. S.; Schieber, N. L.; Templin, R. M.; Steyer, A. M.; Schwab, Y. AMST: Alignment to Median Smoothed Template for Focused Ion Beam Scanning Electron Microscopy Image Stacks. Sci. Rep. 2020, 10 (1), 2004. 10.1038/s41598-020-58736-7.

(21) Krull, A.; Buchholz, T.-O.; Jug, F. Noise2Void - Learning Denoising from Single Noisy Images. arXiv April 5, 2019. 10.48550/arXiv.1811.10980.

(22) Krentzel, D.; Elphick, M.; Domart, M.-C.; Peddie, C. J.; Laine, R. F.; Shand, C.; Henriques, R.; Collinson, L. M.; Jones, M. L. CLEM-Reg: An Automated Point Cloud Based Registration Algorithm for Correlative Light and Volume Electron Microscopy. bioRxiv December 26, 2024, p 2023.05.11.540445. 10.1101/2023.05.11.540445.

(23) Conrad, R.; Narayan, K. Instance Segmentation of Mitochondria in Electron Microscopy Images with a Generalist Deep Learning Model Trained on a Diverse Dataset. Cell Syst. 2023, 14 (1), 58-71.e5. 10.1016/j.cels.2022.12.006.

(24) Archit, A.; Freckmann, L.; Nair, S.; Khalid, N.; Hilt, P.; Rajashekar, V.; Freitag, M.; Teuber, C.; Spitzner, M.; Tapia Contreras, C.; Buckley, G.; von Haaren, S.; Gupta, S.; Grade, M.; Wirth, M.; Schneider, G.; Dengel, A.; Ahmed, S.; Pape, C. Segment Anything for Microscopy. Nat. Methods 2025, 22 (3), 579–591. 10.1038/s41592-024-02580-4.

(25) Gros, O.; Bhickta, C.; Lokaj, G.; Schwab, Y.; Köhler, S.; Banterle, N. Microscopy Nodes: Versatile 3D Microscopy Visualization with Blender. bioRxiv January 14, 2025, p 2025.01.09.632153. 10.1101/2025.01.09.632153.

(26) Dragonfly | 3D Visualization and Analysis Solutions for Scientific and Industrial Data | ORS. https://www.theobjects.com/index.html (accessed 2021-10-07).

(27) Pape, C.; Meechan, K.; Moreva, E.; Schorb, M.; Chiaruttini, N.; Zinchenko, V.; Martinez Vergara, H.; Mizzon, G.; Moore, J.; Arendt, D.; Kreshuk, A.; Schwab, Y.; Tischer, C. MoBIE: A Fiji Plugin for Sharing and Exploration of Multi-Modal Cloud-Hosted Big Image Data. Nat. Methods 2023, 20 (4), 475–476. 10.1038/s41592-023-01776-4.

(28) Perkel, J. M. Python Power-up: New Image Tool Visualizes Complex Data. Nature 2021, 600 (7888), 347–348. 10.1038/d41586-021-03628-7.

(29) Schindelin, J.; Arganda-Carreras, I.; Frise, E.; Kaynig, V.; Longair, M.; Pietzsch, T.; Preibisch, S.; Rueden, C.; Saalfeld, S.; Schmid, B.; Tinevez, J.-Y.; White, D. J.; Hartenstein, V.; Eliceiri, K.; Tomancak, P.; Cardona, A. Fiji: An Open-Source Platform for Biological-Image Analysis. Nat. Methods 2012, 9 (7), 676–682. 10.1038/nmeth.2019.

(30) Asadulina, A.; Conzelmann, M.; Williams, E. A.; Panzera, A.; Jékely, G. Object-Based Representation and Analysis of Light and Electron Microscopic Volume Data Using Blender. BMC Bioinformatics 2015, 16 (1), 229. 10.1186/s12859-015-0652-7.

(31) Gros, A.; Bhickta, C.; Lokaj, G.; Johnston, B.; Schwab, Y.; Köhler, S.; Banterle, N. Microscopy Nodes: Versatile 3D Microscopy Visualization with Blender. EMBO Rep. 2026, 27 (3), 581–597. 10.1038/s44319-025-00654-8.

(32) Labrinidis, A.; Jagadish, H. V. Challenges and Opportunities with Big Data. Proc VLDB Endow 2012, 5 (12), 2032–2033. 10.14778/2367502.2367572.

(33) Jesse, S.; Chi, M.; Belianinov, A.; Beekman, C.; Kalinin, S. V.; Borisevich, A. Y.; Lupini, A. R. Big Data Analytics for Scanning Transmission Electron Microscopy Ptychography. Sci. Rep. 2016, 6 (1), 26348. 10.1038/srep26348.

(34) Vescovi, R.; Li, H.; Kinnison, J.; Keçeli, M.; Salim, M.; Kasthuri, N.; Uram, T. D.; Ferrier, N. Toward an Automated HPC Pipeline for Processing Large Scale Electron Microscopy Data. In 2020 IEEE/ACM 2nd Annual Workshop on Extreme-scale Experiment-in-the-Loop Computing (XLOOP); 2020; pp 16– 22. 10.1109/XLOOP51963.2020.00008.

(35) Munro, I.; García, E.; Yan, M.; Guldbrand, S.; Kumar, S.; Kwakwa, K.; Dunsby, C.; Neil, M. a. a.; French, P. m. w. Accelerating Single Molecule Localization Microscopy through Parallel Processing on a High-Performance Computing Cluster. J. Microsc. 2019, 273 (2), 148–160. 10.1111/jmi.12772.

(36) Hudak, D.; Johnson, D.; Chalker, A.; Nicklas, J.; Franz, E.; Dockendorf, T.; McMichael, B. L. Open OnDemand: A Web-Based Client Portal for HPC Centers. J. Open Source Softw. 2018, 3 (25), 622. 10.21105/joss.00622.

(37) Sun, Y.; Heriche, J.-K.; Kutra, D. The BAND-a Cloud-Based Virtual Desktop for Bioimage Analysis. Zenodo July 9, 2024. https://zenodo.org/records/12699364 (accessed 2025-08-27).

(38) Renton, A. I.; Dao, T. T.; Johnstone, T.; Civier, O.; Sullivan, R. P.; White, D. J.; Lyons, P.; Slade, B. M.; Abbott, D. F.; Amos, T. J.; Bollmann, S.; Botting, A.; Campbell, M. E. J.; Chang, J.; Close, T. G.; Dörig, M.; Eckstein, K.; Egan, G. F.; Evas, S.; Flandin, G.; Garner, K. G.; Garrido, M. I.; Ghosh, S. S.; Grignard, M.; Halchenko, Y. O.; Hannan, A. J.; Heinsfeld, A. S.; Huber, L.; Hughes, M. E.; Kaczmarzyk, J. R.; Kasper, L.; Kuhlmann, L.; Lou, K.; Mantilla-Ramos, Y.-J.; Mattingley, J. B.; Meier, M. L.; Morris, J.; Narayanan, A.; Pestilli, F.; Puce, A.; Ribeiro, F. L.; Rogasch, N. C.; Rorden, C.; Schira, M. M.; Shaw, T. B.; Sowman, P. F.; Spitz, G.; Stewart, A. W.; Ye, X.; Zhu, J. D.; Narayanan, A.; Bollmann, S. Neurodesk: An Accessible, Flexible and Portable Data Analysis Environment for Reproducible Neuroimaging. Nat. Methods 2024, 21 (5), 804–808. 10.1038/s41592-023-02145-x.

(39) Moore, J.; Basurto-Lozada, D.; Besson, S.; Bogovic, J.; Bragantini, J.; Brown, E. M.; Burel, J.-M.; Casas Moreno, X.; de Medeiros, G.; Diel, E. E.; Gault, D.; Ghosh, S. S.; Gold, I.; Halchenko, Y. O.; Hartley, M.; Horsfall, D.; Keller, M. S.; Kittisopikul, M.; Kovacs, G.; Küpcü Yoldaş, A.; Kyoda, K.; le Tournoulx de la Villegeorges, A.; Li, T.; Liberali, P.; Lindner, D.; Linkert, M.; Lüthi, J.; Maitin-Shepard, J.; Manz, T.; Marconato, L.; McCormick, M.; Lange, M.; Mohamed, K.; Moore, W.; Norlin, N.; Ouyang, W.; Özdemir, B.; Palla, G.; Pape, C.; Pelkmans, L.; Pietzsch, T.; Preibisch, S.; Prete, M.; Rzepka, N.; Samee, S.; Schaub, N.; Sidky, H.; Solak, A. C.; Stirling, D. R.; Striebel, J.; Tischer, C.; Toloudis, D.; Virshup, I.; Walczysko, P.; Watson, A. M.; Weisbart, E.; Wong, F.; Yamauchi, K. A.; Bayraktar, O.; Cimini, B. A.; Gehlenborg, N.; Haniffa, M.; Hotaling, N.; Onami, S.; Royer, L. A.; Saalfeld, S.; Stegle, O.; Theis, F. J.; Swedlow, J. R. OME-Zarr: A Cloud-Optimized Bioimaging File Format with International Community Support. Histochem. Cell Biol. 2023, 160 (3), 223–251. 10.1007/s00418-023-02209-1.

(40) Cantoni, M.; Holzer, L. Advances in 3D Focused Ion Beam Tomography. MRS Bull. 2014, 39 (4), 354– 360. 10.1557/mrs.2014.54.

(41) Hashemi, N. S.; Aghdam, R. B.; Ghiasi, A. S. B.; Fatemi, P. Template Matching Advances and Applications in Image Analysis. arXiv October 23, 2016. 10.48550/arXiv.1610.07231.

(42) Tang, Z.; Zhang, Z.; Chen, W.; Yang, W. An SIFT-Based Fast Image Alignment Algorithm for High-Resolution Image. IEEE Access 2023, 11, 42012–42041. 10.1109/ACCESS.2023.3270911.

(43) Anonymous. N2V2 - Fixing Noise2Void Checkerboard Artifacts with Modified Sampling Strategies and a Tweaked Network Architecture; 2022.

(44) Pietzsch, T.; Saalfeld, S.; Preibisch, S.; Tomancak, P. BigDataViewer: Visualization and Processing for Large Image Data Sets. Nat. Methods 2015, 12 (6), 481–483. 10.1038/nmeth.3392.

(45) Ronneberger, O.; Fischer, P.; Brox, T. U-Net: Convolutional Networks for Biomedical Image Segmentation. In Medical Image Computing and Computer-Assisted Intervention – MICCAI 2015; Navab, N., Hornegger, J., Wells, W. M., Frangi, A. F., Eds.; Lecture Notes in Computer Science; Springer International Publishing: Cham, 2015; pp 234–241. 10.1007/978-3-319-24574-4_28.

(46) Morgado, L.; Gómez-de-Mariscal, E.; Heil, H. S.; Henriques, R. The Rise of Data-Driven Microscopy Powered by Machine Learning. arXiv January 10, 2024. 10.48550/arXiv.2401.05282.

(47) Isensee, F.; Rokuss, M.; Krämer, L.; Dinkelacker, S.; Ravindran, A.; Stritzke, F.; Hamm, B.; Wald, T.; Langenberg, M.; Ulrich, C.; Deissler, J.; Floca, R.; Maier-Hein, K. NnInteractive: Redefining 3D Promptable Segmentation. arXiv March 11, 2025. 10.48550/arXiv.2503.08373.

(48) Belevich, I.; Joensuu, M.; Kumar, D.; Vihinen, H.; Jokitalo, E. Microscopy Image Browser: A Platform for Segmentation and Analysis of Multidimensional Datasets. PLOS Biol. 2016, 14 (1), e1002340. 10.1371/journal.pbio.1002340.

(49) Sannerud, R.; Esselens, C.; Ejsmont, P.; Mattera, R.; Rochin, L.; Tharkeshwar, A. K.; De Baets, G.; De Wever, V.; Habets, R.; Baert, V.; Vermeire, W.; Michiels, C.; Groot, A. J.; Wouters, R.; Dillen, K.; Vints, K.; Baatsen, P.; Munck, S.; Derua, R.; Waelkens, E.; Basi, G. S.; Mercken, M.; Vooijs, M.; Bollen, M.; Schymkowitz, J.; Rousseau, F.; Bonifacino, J. S.; Van Niel, G.; De Strooper, B.; Annaert, W. Restricted Location of PSEN2/γ-Secretase Determines Substrate Specificity and Generates an Intracellular Aβ Pool. Cell 2016, 166 (1), 193–208. 10.1016/j.cell.2016.05.020.

(50) Bretou, M.; Sannerud, R.; Escamilla-Ayala, A.; Leroy, T.; Vrancx, C.; Acker, Z. P. V.; Perdok, A.; Vermeire, W.; Vorsters, I.; Keymolen, S. V.; Maxson, M.; Pavie, B.; Wierda, K.; Eskelinen, E.-L.; Annaert, W. Accumulation of APP C-Terminal Fragments Causes Endolysosomal Dysfunction through the Dysregulation of Late Endosome to Lysosome-ER Contact Sites. Dev. Cell 2024, 59 (12), 1571-1592.e9. 10.1016/j.devcel.2024.03.030.

(51) Loginov, S. V.; Fermie, J.; Fokkema, J.; Agronskaia, A. V.; De Heus, C.; Blab, G. A.; Klumperman, J.; Gerritsen, H. C.; Liv, N. Correlative Organelle Microscopy: Fluorescence Guided Volume Electron Microscopy of Intracellular Processes. Front. Cell Dev. Biol. 2022, 10. 10.3389/fcell.2022.829545.

(52) Functional Characterization of Endo-Lysosomal Compartments by Correlative Live-Cell and Volume Electron Microscopy. In Methods in Cell Biology; Academic Press, 2023; Vol. 177, pp 301–326. 10.1016/bs.mcb.2022.12.022.

(53) careamics.github.io/0.1/. https://careamics.github.io/0.1/ (accessed 2025-08-13).

(54) Höck, E.; Buchholz, T.-O.; Brachmann, A.; Jug, F.; Freytag, A. N2V2 -- Fixing Noise2Void Checkerboard Artifacts with Modified Sampling Strategies and a Tweaked Network Architecture. arXiv November 21, 2022. 10.48550/arXiv.2211.08512.

(55) BigDataViewer. https://github.com/bigdataviewer (accessed 2025-08-13).

(56) Gusnard, D.; Kirschner, R. H. Cell and Organelle Shrinkage during Preparation for Scanning Electron Microscopy: Effects of Fixation, Dehydration and Critical Point Drying. J. Microsc. 1977, 110 (1), 51– 57. 10.1111/j.1365-2818.1977.tb00012.x.

(57) Mollenhauer, H. H. Artifacts Caused by Dehydration and Epoxy Embedding in Transmission Electron Microscopy. Microsc. Res. Tech. 1993, 26 (6), 496–512. 10.1002/jemt.1070260604.

(58) Sofroniew, N.; Lambert, T.; Bokota, G.; Nunez-Iglesias, J.; Sobolewski, P.; Sweet, A.; Gaifas, L.; Evans, K.; Burt, A.; Doncila Pop, D.; Yamauchi, K.; Weber Mendonça, M.; Rodríguez-Guerra, J.; Liu, L.; Buckley, G.; Vierdag, W.-M.; Anderson, A.; Monko, T.; Willing, C.; Royer, L.; Can Solak, A.; Harrington, K. I. S.; Abramo, J.; Ahlers, J.; Althviz Moré, D.; Amsalem, O.; Andò, E.; Annex, A.; Aronssohn, C.; Balzaretti, F.; Boone, P.; Bragantini, J.; Bunten, D.; Bussonnier, M.; Caporal, C.; Coccimiglio, I.; Čočková, Z.; Eglinger, J.; Eisenbarth, A.; Freeman, J.; Fukai T. Y.;, Gohlke, C.; Gunalan, K.; Halchenko, Y. O.; Har-Gil, H.; Harfouche, M.; Hilsenstein, V.; Hutchings, K.; Kozar, R.; Lauer, J.; Lichtner, G.; Liu, H.; Liu, Z.; Lowe, A.; Marconato, L.; Martin, S.; McGovern, A.; Migas, L.; Miller, N.; Miñano, S.; Muñoz, H.; Müller, J.-H.; Nauroth-Kreß, C.; Obenhaus, H. A.; Palecek, D.; Pape, C.; Perlman, E.; Theart, R. P.; Pevey, K.; Peña-Castellanos, G.; Pierré, A.; Pinto, D.; Rodríguez-Reza, C. M.; Ross, D.; Russell, C. T.; Ryan, J.; Selzer, G.; Smith, M. B.; Smith, P.; Sofiiuk, K.; Soltwedel, J.; Stansby, D.; Vanaret, J.; Wadhwa, P.; Weigert, M.; Windhager, J.; Winston, P.; Yu, Q.; Zhao, R.; Witz, G. Napari: A Multi-Dimensional Image Viewer for Python. 10.5281/zenodo.18374344.

(59) de Chaumont, F.; Dallongeville, S.; Chenouard, N.; Hervé, N.; Pop, S.; Provoost, T.; Meas-Yedid, V.; Pankajakshan, P.; Lecomte, T.; Le Montagner, Y.; Lagache, T.; Dufour, A.; Olivo-Marin, J.-C. Icy: An Open Bioimage Informatics Platform for Extended Reproducible Research. Nat. Methods 2012, 9 (7), 690–696. 10.1038/nmeth.2075.

